# Altered aperiodic EEG spectral power during speech perception task is associated with verbal communication in youths with Autism Spectrum Disorder

**DOI:** 10.64898/2025.12.29.696902

**Authors:** Vardan Arutiunian, Megha Santhosh, Emily Neuhaus, Heather Borland, Raphael A. Bernier, Susan Y. Bookheimer, Mirella Dapretto, Abha R. Gupta, Allison Jack, Shafali Jeste, James C. McPartland, Adam Naples, John D. Van Horn, Kevin A. Pelphrey, Sara J. Webb

## Abstract

Most children with Autism Spectrum Disorder (ASD) have co-occurring language impairment, but its neural mechanisms are not well known. Excitation (E) / inhibition (I) imbalance is considered as a key neurobiological mechanism of ASD, and several electroencephalography (EEG)-based E/I balance metrics have been proposed in the previous studies. The goal of the present research was to focus on these metrics abstracted from the speech perception task to investigate their relation to language/communication in autistic youths. We used a high-density 128-channel EEG to register neural responses during speech perception task in the sex- and age-matched groups of youths with ASD (*N* = 162) and typically developing (TD) controls (*N* = 144), aged 7–18 years old. The results revealed alterations in the E/I measures in the ASD group, pointing to a higher level of excitation or neural ‘noise’ in the cortex as well as broadband reduction of spectral power during speech perception. A greater neural ‘noise’ reflected in the reduction of aperiodic exponent and offset was associated with lower verbal communication skills in youths with ASD. The findings suggested that the higher ‘noisiness’ in the cortical systems may be a relevant marker to monitor in relation to language/communication in ASD.

## Introduction

Autism Spectrum Disorder (ASD) is a neurodevelopmental condition associated with the difficulties in social interaction / communication and restricted and repetitive behaviors (1). It is known that there is a high heterogeneity in language abilities in individuals with ASD, from above average language skills (∼25%) to severe language impairments with minimal or even no verbal skills (2–5). Although language impairment is not a diagnostic criterion of ASD, previous studies have shown that difficulties in language / verbal communication are one of the first signs of ASD at early stages of development and can be related to later ASD outcomes (6,7). However, neural mechanisms of these difficulties are still unknown, and there is a need to develop a set of biomarkers of language / verbal communication impairments in ASD for using as objective measures in clinical trial studies.

Excitation (E) / inhibition (I) imbalance in different brain networks is considered an important neurobiological mechanism in ASD (8–13). At the system level, multiple studies have suggested that altered interplay between glutamatergic (excitatory) and gamma-aminobutyric acidergic (GABAergic; inhibitory) neurotransmission causes a reduction in signal-to-noise ratio in key neural circuits (12). This, in turn, may cause an enhanced (or reduced) “noise” in the cortex, which impacts synaptic plasticity during development and results in less effective information processing (12,14,15). Although alterations in E/I balance can have different causes (elevated E over I as well as reduced E over I, see [8,16–18]), most studies reported increased excitatory activity in the autistic brain (11,12). Several electro-/magnetoencephalography (EEG/MEG)-based E/I balance biomarkers have been proposed to index language and verbal communication impairments in ASD and related disorders (19–25).

Gamma-band neural activity (30–80Hz) in EEG / MEG is one of the most frequently reported non-invasive measures of E/I balance in the cortex, and it is generated mostly by GABAergic interneurons, expressing calcium-binding protein parvalbumin (PV+ inhibitory cells) (12,26–31). Previous studies have shown atypicalities in gamma activity and relationships between gamma power and language / verbal communication skills in autistic individuals and their first-degree relatives (19,21,32–41). For example, atypical gamma activity was detected in response to low-level auditory stimuli, such as 40Hz amplitude-modulated tones and sweeps, and those atypicalities were associated with lower language skills (19,36). Similar abnormalities in gamma activity were detected in autistic individuals and their first-degree relatives when presenting linguistic stimuli, such as syllables, words, and sentences (21,33,34). In addition, resting-state EEG gamma oscillations were also associated with language skills in toddlers with ASD and their siblings and/or parents as well as in individuals with Fragile X Syndrome, a single-gene disorder related to ASD (25,41,42).

Development of newer methods for EEG/MEG power analysis has allowed parametrization of spectral power into periodic and aperiodic components to provide more accurate estimation of both oscillatory and non-oscillatory (broadband) activity associated with different neurobiological mechanisms (43–45). The aperiodic component of the signal can be defined by the 1/f power law distribution that underlies the absolute power spectra (43,45–48) and consists of aperiodic exponent and aperiodic offset. Different modeling studies have shown that the exponent reflects E/I balance and its characteristics can indicate whether the circuit has increased excitatory or inhibitory activity (46), while offset reflects broadband neuronal firing (47). Investigation of the aperiodic signal together with the oscillatory part of the spectra may significantly contribute to our understanding of the mechanisms at the system level (24).

Although this approach is relatively novel, there is evidence of a relationship between the aperiodic component and language/verbal communication skills in ASD and related conditions. For example, Wilkinson and Nelson (25) found that the aperiodic component of spectral power in the gamma frequency range was altered in Fragile X Syndrome and was related to communication skills in this population. Using MEG, Arutiunian and colleagues (21) demonstrated that aperiodic offset in the auditory cortex during pre-stimulus/baseline period was associated with language abilities in children with ASD. Finally, Wilkinson et al. (24) observed that developmental changes in aperiodic offset and exponent (from 3 to 12 months) predict lower language skills at 18 months in infants with a family history of ASD.

To summarize, previous studies using different EEG/MEG measures of E/I balance have demonstrated its alterations in ASD. Findings have largely suggested elevated excitatory activity in the autistic brain in response to auditory and speech stimuli, with increased excitation related to lower language and/or communication skills. The aims of the present study were to: 1) comprehensively investigate these neural measures in response to speech stimuli in a large, sex-balanced group of children with and without ASD, and 2) explore associations between these measures and language/communication in autistic youth. *First,* for between-group comparisons we applied a parameterization approach to calculate periodic and aperiodic components of power spectra. We abstracted measures related to E/I balance used in the previous studies: oscillatory gamma power, aperiodic exponent, and aperiodic offset. Based on findings from previous studies, we hypothesized that E/I balance would be altered in ASD, with evidence of elevated excitation. *Second,* we hypothesized that this elevation could potentially be related to language/communication in the ASD group.

## Methods

### Participants

In total, 306 English-speaking participants (7 to 18 years) took part in the study: 162 youth with ASD (90 male, 72 female) and 144 age- and sex-matched TD controls (77 male, 67 female). Sex was based on parent report of sex assigned at birth. Data were collected from four sites as a part of the GENDAAR Autism Center for Excellence Network, including Seattle Children’s Research Institute, Boston Children’s Hospital, the University of California in Los Angeles, and Yale University.

The study was approved by the Yale Institutional Review Board, the UCLA Office of Human Research Protection Program, Boston Children’s Hospital Institutional Review Board, USC Office for the Protection of Research Subjects, and the University of Virginia Institutional Review Board for Health Sciences Research. All performed procedures were in accordance with the Declaration of Helsinki. All minor children provided verbal assent to participate in the study and were informed that they can withdraw from the study at any time. A written consent form was obtained from a parent of each child participating in the study.

### Behavioral assessment

All participants from the ASD group had a clinical diagnosis based on the DSM-4–TR or DSM-5 (1,49) and the diagnosis was confirmed with the Autism Diagnosis Observation Schedule – Second Edition (50), Autism Diagnostic Interview – Revised (51), caregiver-reported developmental history, and expert clinical judgment. Adaptive skills of all youths were measured with the Vineland Adaptive Behavior Scales – Second Edition (VABS-2), and standard scores (SS) in communication, socialization, and daily living skills domains were calculated with higher scores reflecting better adaptive skills (52). Verbal and nonverbal IQ were estimated based on the Differential Ability Scales – Second Edition (DAS-II) School Aged Cognitive Battery (53). Language abilities were measured with the Clinical Evaluation of Language Fundamentals – Fourth Edition (CELF-4), a standardized assessment tool that covers basic structural language skills at different linguistic levels (vocabulary, morphosyntax, semantics, and pragmatics) in both production and comprehension (54). Participants were administered only those subtests necessary to calculate the CELF-4 Core Language SS, which was used as a measure of overall language skills. Parents were asked to report on their child’s autistic traits and social skills using the Social Responsiveness Scale – Second Edition (SRS-2); T-scores for social cognition, social communication, and restricted and repetitive behaviors were calculated for all youths with the higher scores reflecting greater severity (55). Descriptive statistics are presented in Table 1.

**Table 1.** Behavioral characteristics of participants, M(SD). Significance is Significance is labeled with *\*p* < 0.05, *\*\*p* < 0.01, *\*\*\*p* < 0.001 and highlighted in bold.

| Characteristics | Group |  | Statistics |
| --- | --- | --- | --- |
| | ASD ($N = 162$) | TD ($N = 144$) | |
| Age, months | 151.2 (34.9) | 156.5 (35.3) | $t = -1.32, p = 0.18$ |
| Sex, $N$ of female participants | 72 | 67 | $\chi^2(1) = 0.13, p = 0.71$ |
| CELF-4 Core Language SS | 91.43 (20.90) | 110.62 (11.26) | $t = -8.74, p < \mathbf{0.001^{***}}$ |
| Vineland-2 |  |  |  |
| *Communication SS* | 76.1 (11.8) | 99.3 (13.2) | $t = -15.9, p < \mathbf{0.001^{***}}$ |
| *Socialization SS* | 72.6 (12.1) | 101.2 (12.6) | $t = -19.9, p < \mathbf{0.001^{***}}$ |
| *Daily living skills SS* | 76.4 (13.7) | 97.7 (14.2) | $t = -13.1, p < \mathbf{0.001^{***}}$ |
| Verbal IQ | 100.6 (20.2) | 113.1 (15.6) | $t = -6.0, p < \mathbf{0.001^{***}}$ |
| Nonverbal IQ | 100.2 (17.5) | 109.1 (15.1) | $t = -4.7, p < \mathbf{0.001^{***}}$ |
| SRS-2 T-score |  |  |  |
| *Social cognition* | 71.6 (10.9) | 43.8 (5.2) | $t = 27.9, p < \mathbf{0.001^{***}}$ |
| *Social communication* | 74.5 (12.0) | 43.9 (5.2) | $t = 28.2, p < \mathbf{0.001^{***}}$ |
| *Restricted and repetitive behavior* | 73.7 (12.6) | 44.4 (3.5) | $t = 27.0, p < \mathbf{0.001^{***}}$ |

Exclusion criteria were twin status, history and/or presence of known chromosomal syndromes/single-gene conditions related to autism (e.g., Fragile X Syndrome), co-occurring neurological conditions (e.g., epilepsy), significant visual and auditory impairments, or sensory-motor difficulties that would prevent completion of study procedures. TD children had no first or second degree family members with ASD, and no elevation of autism traits according to parent report on the SRS-2 (total T-score < 60) or the Social Communication Questionnaire (56), raw score < 11.

### Experimental paradigm and procedure

We used the implicit word segmentation paradigm described in detail in (21) but see also (57–62). In brief, the experiment consisted of two phases. During the first (exposure) phase, youths heard three-syllabic pseudowords generated from the set of 12 different phonemes (e.g., *pa-bi-ku*), resulting in 180 exposures over ∼2.5 minutes. The second (test) phase consisted of 96 trials; duration was 2 minutes 16 seconds. Analyses were restricted to the second (test) phase of the experiment. Each trial consisted of a three-syllabic pseudoword with an average duration of ∼900 ms, followed by a 500–750 ms intertrial interval. A random half of these trials (*N* = 48) used the same pseudowords presented in the first phase during exposure (e.g., *pa-bi-ku*), i.e., ‘familiar’ items. The remaining random trials (*N* = 48) were constructed by combining the last syllable of each familiar pseudoword with the first two phonemes of other pseudowords (e.g., *pa-bi-ku* and *go-la-tu* became *ku-go-la* and *tu-pa-bi*), i.e., ‘unfamiliar’ items. The auditory stimuli were presented using a speaker (Logitech speaker system X320) with the same loudness across all participants and sites (65 dB). All participants were instructed to look at the screen and to listen carefully to the ‘robot language’. A static robot was presented on the screen simultaneously with the auditory stimuli during the experiment.

### EEG data acquisition and processing

At all four sites, EEG data were collected with EGI 128-channel Net Amps 300 system with HydroCel nets (Magstim EGI Inc., Eugene OR), using Net Station 4.4.2, 4.5.1, or 4.5.2 with a standard Net Station acquisition template. Nets were available without outriders (eye electrodes 125, 126, 127, and 128) for participants with facial sensory sensitivities. The participant’s behavior was video recorded during EEG collection. Data were collected at 500 Hz sampling rate, referenced to Cz electrode (vertex), and impedances were < 50 kΩ.

To calculate power spectral density (PSD) we used the Batch EEG Automated Processing Platform, BEAPP (63) in MATLAB 2021a, consisting of following steps: (1) formatting the MFF file for Matlab; (2) band-pass filter 1–100 Hz; (3) down sampling from 500 Hz to 250 Hz; (4) implementation of the Harvard Automated Preprocessing Pipeline for EEG (HAPPE) module for artifact detection and rejection (64), including removal of 60 Hz line noise, rejection of bad channels, wavelet enhanced thresholding, Independent Component Analysis (ICA) with automated component rejection, bad channel interpolation, and re-referencing to average; (5) segmentation of the continuous file into 1 second epochs (each epoch consisted of one three-syllabic pseudoword); (6) rejection of bad segments (± 40 μV); (7) calculation of the PSD using Hanning window on clean segments.

### EEG variables calculation

To model the periodic and aperiodic components of PSD (43), we used the *specparam* toolbox (45) in Python v3.10 with the following settings: *peak_width_limit* = [1.5, 8], *n_peaks* = 6, *peak_height* = 0.10, *peak_threshold* = 2, and *frequency_range* = [2,55]. For each participant, the model provided two parameters (aperiodic offset and aperiodic exponent) to describe the aperiodic 1/f signal. To calculate periodic power (or aperiodic-adjusted power), the aperiodic signal from the model was subtracted from the raw power spectrum, resulting in a flattened spectrum. Three measures were used for statistical analysis: aperiodic exponent, aperiodic offset, and periodic gamma power (35–54.99Hz). The measures were calculated for nine regions of interest (ROIs; see Figure 1A).

**Figure 1.**
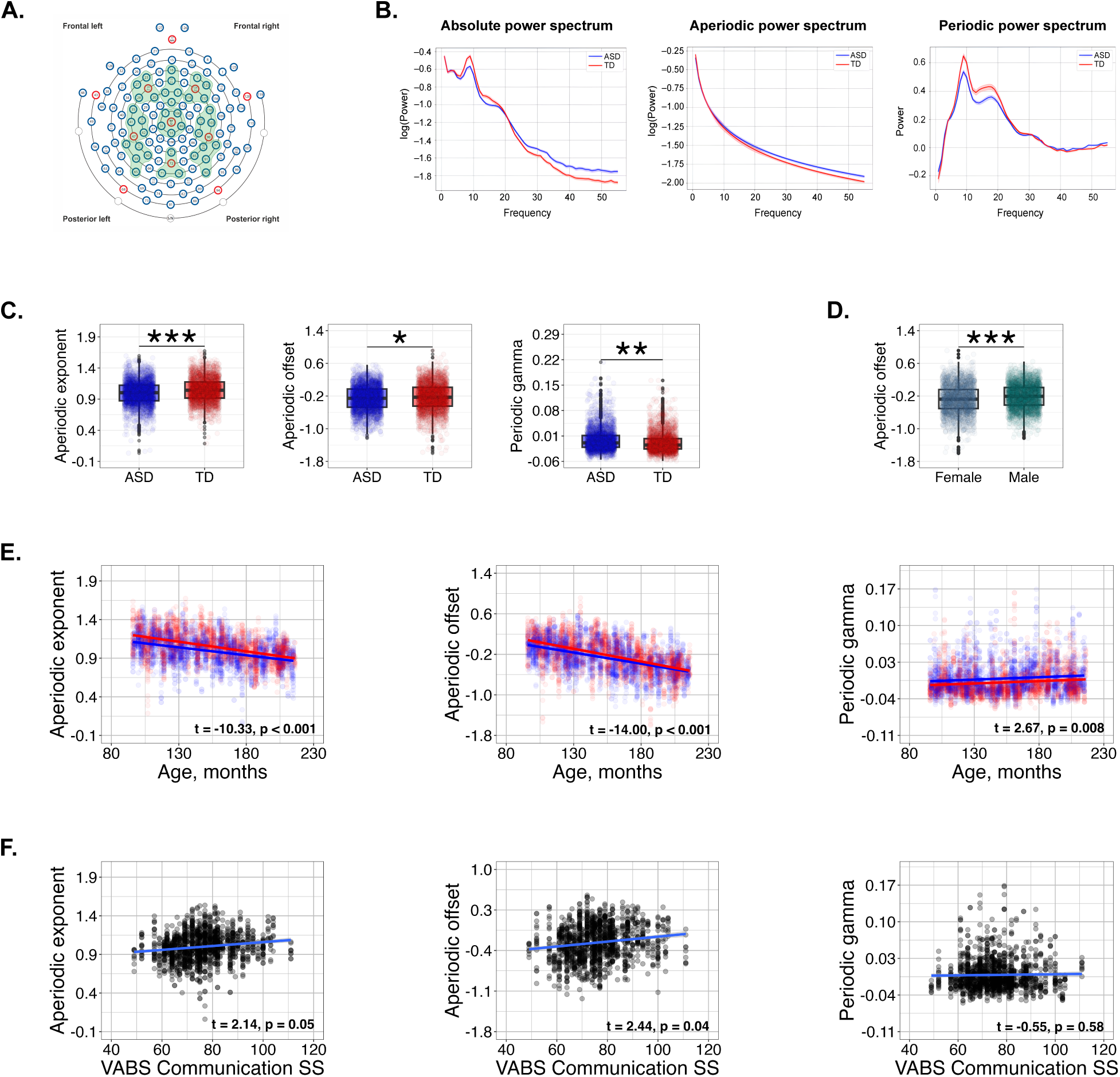
EEG neural measures in response to speech stimuli in youths with and without Autism Spectrum Disorder: **A.** EEG montage with channels indicated. Channel numbers for regions are frontal left (20, 23, 24, 27, 28), frontal midline (5, 6, 11, 12, 16), frontal right (3, 117, 118, 123, 124), central left (35, 36, 41, 42, 47), central midline (7, 31, 55, 80, 106), central right (93, 98, 103, 104, 110), posterior left (51, 52, 59, 60, 65), posterior midline (62, 71, 72, 76), (9) posterior right (85, 90, 91, 92, 97). **B.** Absolute, aperiodic and periodic power spectral density (for central left ROI ‘familiar’ items, for visualization purposes). **C.** Boxplots representing between-group differences in neural measures. **D.** Boxplots representing sex differences in aperiodic offset. **E.** Relationships between EEG neural responses and age. **F.** Relationships between EEG neural responses and verbal communication. ASD = Autism Spectrum Disorder, TD = typically developing, VABS = Vineland Adaptive Behavior Scales. Significance is labeled with *\*p* < 0.05, *\*\*p* < 0.01, *\*\*\*p* < 0.001; *p*-values are FDR-corrected.

The ASD and TD groups did not differ in the number of artifact-free epochs in any condition: ‘familiar’ items or pseudowords, *M*_ASD_ = 43.2(1.9) vs. *M*_TD_ = 43.2(3.6), *t*(217.1) = – 0.21, *p* = 0.82; ‘unfamiliar’ items, *M*_ASD_ = 43.0(2.0) vs. *M*_TD_ = 43.4(3.3), *t*(228.6) = –1.37, *p* = 0.17.

### Statistical analysis

Statistical analysis was performed in R (65) with the *lme4* package (66). The tables for model outcomes (Tables 2–4) were created with the *sjPlot* package (67), data were plotted with *ggplot2* (68), and the figures, representing neural responses, were created with python data visualization library *matplotlib* (69). The structure of the models will be specified further in the Results section. A correction for multiple comparisons (false discovery rate, FDR) was applied to the models, and p-values for significant predictors were corrected with *p.adjust.method* in R.

**Table 2.** The outputs of the models. Significance is labeled with *\*p* < 0.05, *\*\*p* < 0.01, *\*\*\*p* < 0.001 and highlighted in bold. All significant *p*-values are FDR-corrected.

| Predictors | Aperiodic exponent |  |  |  | Aperiodic offset |  |  |  | Periodic gamma power |  |  |  |
| --- | --- | --- | --- | --- | --- | --- | --- | --- | --- | --- | --- | --- |
| | $\beta$ | SE | $t$ | $p$ | $\beta$ | SE | $t$ | $p$ | $\beta$ | SE | $t$ | $P$ |
| (Intercept) | 1.36 | 0.04 | 36.67 | **<0.001\*\*\*** | 0.58 | 0.06 | 10.11 | **<0.001\*\*\*** | -0.02 | 0.01 | -2.91 | **0.004\*\*** |
| Group, ASD | -0.06 | 0.02 | -3.95 | **<0.001\*\*\*** | -0.06 | 0.02 | -2.39 | **0.017\*** | 0.01 | 0.00 | 2.98 | **0.005\*\*** |
| Condition, unfamiliar | -0.00 | 0.00 | -0.01 | 0.989 | 0.00 | 0.01 | 0.31 | 0.759 | -0.00 | 0.00 | -0.36 | 0.721 |
| Sex, male | 0.02 | 0.02 | 1.53 | 0.127 | 0.10 | 0.02 | 4.00 | **<0.001\*\*\*** | -0.00 | 0.00 | -1.84 | 0.099 |
| Region, central | 0.03 | 0.00 | 7.49 | **<0.001\*\*\*** | -0.18 | 0.01 | -34.79 | **<0.001\*\*\*** | -0.01 | 0.00 | -8.65 | **<0.001\*\*\*** |
| Region, parietal | 0.00 | 0.00 | 0.84 | 0.399 | -0.03 | 0.01 | -5.45 | **<0.001\*\*\*** | 0.00 | 0.00 | 1.78 | 0.112 |
| Laterality, midline | 0.08 | 0.00 | 23.28 | **<0.001\*\*\*** | -0.05 | 0.01 | -10.03 | **<0.001\*\*\*** | -0.00 | 0.00 | -5.40 | **<0.001\*\*\*** |
| Laterality, right | 0.00 | 0.00 | 1.04 | 0.298 | -0.05 | 0.01 | -9.18 | **<0.001\*\*\*** | -0.00 | 0.00 | -4.24 | **<0.001\*\*\*** |
| Age | -0.00 | 0.00 | -10.33 | **<0.001\*\*\*** | -0.00 | 0.00 | -14.00 | **<0.001\*\*\*** | 0.00 | 0.00 | 2.67 | **0.008\*\*** |
| Group × Condition | -0.00 | 0.01 | -0.21 | 0.837 | -0.01 | 0.01 | -0.95 | 0.341 | 0.00 | 0.00 | 0.27 | 0.790 |
| **Random effects** |  |  |  |  |  |  |  |  |  |  |  |  |
| $\sigma^2$ | | | 0.01 | | | | 0.02 | | | | 0.00 | |
| $\tau_{00}$ ID | | | 0.02 | | | | 0.04 | | | | 0.00 | |
| ICC |  |  | 0.62 |  |  |  | 0.64 |  |  |  | 0.56 |  |
| $N_{ID}$ | | | 306 | | | | 306 | | | | 306 | |
| Observations |  |  | 5508 |  |  |  | 5508 |  |  |  | 5508 |  |
| **Marginal $R^2$ /** |  |  |  |  |  |  |  |  |  |  |  |  |
| Conditional $R^2$ | | | 0.230/0.709 | | | | 0.353/0.767 | | | | 0.045/0.582 | |

**Table 3.** Relationships between EEG neural measures and verbal communication in youths with ASD. The outputs of the models. Significance is labeled with *\*p* < 0.05, *\*\*p* < 0.01, *\*\*\*p* < 0.001 and highlighted in bold. All significant *p*-values are FDR-corrected.

| Predictors | Aperiodic exponent |  |  |  | Aperiodic offset |  |  |  | Periodic gamma power |  |  |  |
| --- | --- | --- | --- | --- | --- | --- | --- | --- | --- | --- | --- | --- |
| | $\beta$ | $SE$ | $t$ | $p$ | $\beta$ | $SE$ | $t$ | $p$ | $\beta$ | $SE$ | $t$ | $p$ |
| (Intercept) | 0.75 | 0.12 | 6.35 | **<0.001\*\*\*** | -0.74 | 0.20 | -3.75 | **<0.001\*\*\*** | 0.01 | 0.02 | 0.53 | 0.598 |
| VABS Communication | 0.00 | 0.00 | 2.14 | **0.05\*** | 0.01 | 0.00 | 2.44 | **0.04\*** | -0.00 | 0.00 | -0.55 | 0.583 |
| Sex, male | 0.14 | 0.16 | 0.84 | 0.402 | 0.29 | 0.27 | 1.09 | 0.275 | -0.02 | 0.02 | -0.82 | 0.414 |
| VABS Communication × Sex | 0.00 | 0.00 | -0.72 | 0.473 | -0.00 | 0.00 | -0.88 | 0.381 | 0.00 | 0.00 | 0.71 | 0.477 |
| **Random effects** |  |  |  |  |  |  |  |  |  |  |  |  |
| $\sigma^2$ | | | 0.01 | | | | 0.02 | | | | 0.00 | |
| T<sub>00</sub> ID |  |  | 0.02 |  |  |  | 0.06 |  |  |  | 0.00 |  |
| T<sub>00</sub> region |  |  | 0.00 |  |  |  | 0.01 |  |  |  | 0.00 |  |
| T<sub>00</sub> laterality |  |  | 0.00 |  |  |  | 0.00 |  |  |  | 0.00 |  |
| ICC |  |  | 0.70 |  |  |  | 0.76 |  |  |  | 0.55 |  |
| N<sub>ID</sub> |  |  | 158 |  |  |  | 158 |  |  |  | 158 |  |
| N<sub>region</sub> |  |  | 3 |  |  |  | 3 |  |  |  | 3 |  |
| N<sub>laterality</sub> |  |  | 3 |  |  |  | 3 |  |  |  | 3 |  |
| Observations |  |  | 1422 |  |  |  | 1422 |  |  |  | 1422 |  |
| Marginal R<sup>2</sup> / Conditional R<sup>2</sup> |  |  | 0.025/0.708 |  |  |  | 0.036/0.772 |  |  |  | 0.004/0.553 |  |

**Table 4.** Relationships between EEG neural measures and language skills in youths with ASD. The outputs of the models. Significance is labeled with *\*p* < 0.05, *\*\*p* < 0.01, *\*\*\*p* < 0.001 and highlighted in bold. All significant *p*-values are FDR-corrected.

| Predictors | Aperiodic exponent |  |  |  | Aperiodic offset |  |  |  | Periodic gamma power |  |  |  |
| --- | --- | --- | --- | --- | --- | --- | --- | --- | --- | --- | --- | --- |
| | $\beta$ | SE | $t$ | $p$ | $\beta$ | SE | $t$ | $p$ | $\beta$ | SE | $t$ | $p$ |
| (Intercept) | 1.02 | 0.11 | 9.42 | **<0.001\*\*\*** | -0.25 | 0.18 | -1.45 | 0.147 | -0.01 | 0.02 | -0.38 | 0.703 |
| CELF | -0.00 | 0.00 | -0.29 | 0.773 | -0.00 | 0.00 | -0.20 | 0.00 | 0.00 | 0.38 | 0.701 | 0.00 |
| Sex, male | -0.02 | 0.14 | -0.16 | 0.876 | -0.01 | 0.22 | -0.01 | 0.02 | -0.53 | 0.598 | -0.01 | 0.02 |
| CELF × Sex | 0.00 | 0.00 | 0.31 | 0.755 | 0.00 | 0.00 | 0.00 | 0.00 | 0.43 | 0.670 | 0.00 | 0.00 |
| **Random effects** |  |  |  |  |  |  |  |  |  |  |  |  |
| $\sigma^2$ | | | 0.01 | | | | 0.02 | | | | 0.00 | |
| T<sub>00</sub> ID |  |  | 0.02 |  |  |  | 0.06 |  |  |  | 0.00 |  |
| T<sub>00</sub> region |  |  | 0.00 |  |  |  | 0.01 |  |  |  | 0.00 |  |
| T<sub>00</sub> laterality |  |  | 0.00 |  |  |  | 0.00 |  |  |  | 0.00 |  |
| ICC |  |  | 0.69 |  |  |  | 0.75 |  |  |  | 0.53 |  |
| N<sub>ID</sub> |  |  | 118 |  |  |  | 118 |  |  |  | 118 |  |
| N<sub>region</sub> |  |  | 3 |  |  |  | 3 |  |  |  | 3 |  |
| N<sub>laterality</sub> |  |  | 3 |  |  |  | 3 |  |  |  | 3 |  |
| Observations |  |  | 1062 |  |  |  | 1062 |  |  |  | 1062 |  |
| Marginal R<sup>2</sup> / |  |  |  |  |  |  |  |  |  |  |  |  |
| Conditional |  |  | 0.003/0.694 |  |  |  | 0.015/0.759 |  |  |  | 0.008/0.532 |  |
| R<sup>2</sup> |  |  |  |  |  |  |  |  |  |  |  |  |

### Data availability

The EEG and behavioral data from the current study are available via the National Institute of Mental Health Data Archive Data Collection #2021.

## Results

### Sample characterization

As expected, youths with ASD differed from the TD group in all behavioral measures, having lower skills in language / communication and non-verbal cognition as well as higher presence of autistic traits (see Table 1). There was no difference in age and sex between groups.

### Group, condition, sex differences and age-related changes in neural measures

Figure 1B represents the example of absolute, aperiodic, and periodic spectral power in response to speech stimuli for one ROI (central left). In order to investigate between-group differences in EEG neural measures in response to speech stimuli, we fitted linear mixed-effect models with the EEG metrics as dependent variables and included the main effects of group (‘TD’ as a reference), condition (‘familiar’ as a reference), sex (‘female’ as a reference), region (‘frontal’ as a reference), laterality (‘left’ as a reference), age, and group ×· condition interaction as fixed effects and participants as random intercept. The structure of the models was as follows: *lmer(EEG metric ∼ group* × *condition + sex + region + laterality + age + (1 | ID), data = data, control = lmerControl(optimizer = “Nelder_Mead”))*.

#### Region of interest (ROI) and laterality effects in EEG neural measures

We indicated significant region and laterality effects for all neural measures (see Table 2 for summary).

Specifically, when comparing to frontal ROI, aperiodic exponent was significantly higher in central ROI, *β* = 0.03, *t* = 7.49, FDR-corrected *p* < 0.001; aperiodic offset was significantly lower in central and parietal ROIs, *β* = –0.18, *t* = –34.79, FDR-corrected *p* < 0.001 and *β* = – 0.03, *t* = –5.45, FDR-corrected *p* < 0.001; and periodic gamma power was significantly lower in central ROI, *β* = –0.01, *t* = –8.65, FDR-corrected *p* < 0.001.

Regarding laterality effects, when comparing to the left EEG subchannels, aperiodic exponent was significantly higher in the midline area, *β* = 0.08, *t* = 23.28, FDR-corrected *p* < 0.001; aperiodic offset was significantly lower in the midline and right areas, *β* = –0.05, *t* = – 10.03, FDR-corrected *p* < 0.001 and *β* = –0.05, *t* = –9.18, FDR-corrected *p* < 0.001; and periodic gamma power was significantly lower the midline and right areas, *β* = –0.00, *t* = –5.40, FDR-corrected *p* < 0.001 and *β* = –0.00, *t* = –4.24, FDR-corrected *p* < 0.001.

#### Group, condition, and sex effects in EEG neural measures

Between-group comparisons revealed significant differences in all the three EEG neural metrics in response to speech stimuli (see Table 2, Figure1C). Autistic youths had reduced aperiodic exponent, *β* = –0.06, *t* = 5.46, FDR-corrected *p* < 0.001; reduced aperiodic offset, *β* = –0.06, *t* = 2.39, FDR-corrected *p* = 0.017; and elevated periodic gamma power, *β* = 0.01, *t* = 2.98, FDR-corrected *p* = 0.005.

Condition as well as group × condition effects were not significant for all EEG measures (see Table 2).

In male participants in comparison to females, it was observed a significantly higher aperiodic offset, *β* = 0.10, *t* = 4.00, FDR-corrected *p* < 0.001 (Table 2, Figure1D).

#### Age-related changes of EEG neural measures

Linear age-related changes were revealed in the strength of all neural measures (see Table 2, Figure 1E): decrease in aperiodic exponent, *β* = –2.29×10^-3^, *t* = –10.33, FDR-corrected *p* < 0.001; decrease in aperiodic offset, *β* = –4.82×10^-3^, *t* = –14.00, FDR-corrected *p* < 0.001; and increase in periodic gamma power, *β* = 9.45×10^-5^, *t* = 2.67, FDR-corrected *p* = 0.008. The mean trajectories demonstrated similar age-related changes in neural measures in both groups of youths.

### The relationships between neural measures and clinical phenotype in youths with ASD

This analysis aimed to investigate whether the EEG measures in response to speech stimuli were associated with language and communication skills in the ASD group when considering a sex effect. Six linear mixed-effects models with EEG metrics as dependent variables, behavioral measures as main effects (CELF Core Language SS, VABS Communication SS) and behavioral measure × sex interaction as fixed effects were fitted. As we did not have a-priory hypothesis on region/laterality, we included it as a random intercepts (with the random effect of participants) in the models. The structure of the models was as follows: *lmer(EEG metric ∼ behavioral measure* × *sex + (1 | ID) + (1 | region) + (1 | laterality), data = data, control = lmerControl(optimizer = “Nelder_Mead”)).* As there was a very high correlation between ‘familiar’ and ‘unfamiliar’ items for all neural measures, brain-behavior relationship analysis was performed only for ‘unfamiliar’ items to avoid familiarity/habituation effect: ASD group, aperiodic exponent, *r* = 0.91, *p* < 0.001; aperiodic offset, *r* = 0.95, *p* < 0.001; periodic gamma power, *r* = 0.83, *p* < 0.001.

The output of the models can be seen in Table 3. The results showed that a reduction of the aperiodic components (both exponent and offset) of the EEG spectral power during speech perception was associated with lower verbal communication skills in ASD: aperiodic exponent, *β* = 0.00, SE = 0.00, *t* = 2.14, FDR-corrected *p* = 0.05; aperiodic offset, *β* = 0.01, SE = 0.00, *t* = 2.44, FDR-corrected *p* = 0.04; no relationship was detected for periodic gamma power, *β* = – 0.00, SE = 0.00, *t* = –0.55, FDR-corrected *p* = 0.58 (Figure1F, Table 3).

No significant relationships were detected between CELF Core Language SS and neural responses (Table 4). The results did not reveal any interaction with the sex (Table 3, 4).

## Discussion

The present study investigated neural mechanisms of speech perception in a large sample of youths with ASD and relation of these mechanisms to behavioral measures of language and verbal communication. Three potential EEG E/I balance measures have been abstracted from the EEG during speech perception task: aperiodic exponent, aperiodic offset, and periodic gamma power. In general, we revealed altered E/I balance in the ASD group compared to TD controls, and variability in these E/I measures was associated with verbal communication skills in youths with ASD.

Between-group comparisons of EEG E/I balance measures during speech perception task demonstrated significant differences all three variables: autistic youths had reduced aperiodic exponent and aperiodic offset and elevated periodic gamma power. Although those are separate neural measures abstracted from the EEG signal, all between-group differences had similar functional direction: youths with ASD had increased excitation in comparison to the TD group. Although we used updated parameterization methods, our results are consistent with the previous studies (11,12,14,15).

Aperiodic exponent, one of the primary measures of E/I balance, was significantly reduced in the ASD group, and research in humans and animals as well as computational modeling have revealed that this reflects elevated excitatory activity in the cortex and, subsequently, elevated neural ‘noise’ in the brain networks (43,45,46). Remarkably, the ASD group had also a reduced aperiodic offset which is associated with a broadband neural firing and population spiking (47,48). These two EEG measures are tightly related to each other. At the system level, greater excitation in the neural circuits (reduced exponent) can lead to a reduction of broadband neural activity (thus, to a higher ‘noisiness’ of the cortex), so that higher E/I ratio is related to greater randomness in the patters of neural firing (70–74). Likewise, elevated periodic gamma power in youths with ASD reflected the same atypical functioning of neural populations as the aperiodic component of neural spectra with the evidence of increased E/I ratio. This is consistent with previous studies that have reported both increased gamma power (21,25) in response to auditory and language stimuli in individuals with ASD.

Although we did not conduct source estimation, we hypothesized that the main generators of neural activity were primary and secondary temporal regions in the left and right hemispheres given the nature of the task (perception of speech stimuli presented auditorily). This corresponds to previous studies that showed alterations in gamma power and aperiodic component of the spectra in the auditory cortex of autistic individuals (19–21,33,34,36). All of these alterations observed in the ASD group (e.g., increased excitation and broadband reduction of neural firing leading to increased random neural ‘noise’ in the circuit) can affect the precise timing and coordination of neural activity, which is essential for information processing (71,73). Thus, this alteration in neural activity can impact temporal cortex ‘functions’ in youths with ASD, including not only language and verbal communication but also more general social communication (75,76).

Indeed, we found relationships between atypical neural activity in response to speech stimuli and behavioral measures in youths with ASD. Reduction of both aperiodic exponent and aperiodic offset was related to lower communication skills in autistic individuals; this is consistent with the limited number of existing studies on aperiodic neural activity in neurodevelopmental disorders (21,24,25). When perceiving speech (three-syllabic pseudowords in this study), the temporal cortex supports different hierarchical mechanisms for sound processing, such as decomposition of complex sound with specific temporal and frequency features into pure tones by primary auditory cortex, coding temporal fine units, analyzing the properties of a sound in the short temporal windows, integration of those features into complex signal by secondary auditory cortex, etc. (77–83). To process this information, very precise neural computations are needed, and it has been shown in mice that deregulation of these computations by increasing neural ‘noise’ and shifting E/I balance towards more excitation leads to sound perception and recognition disorders (84). In our ASD group, we revealed a reduction of aperiodic exponent and offset in speech perception task, which is consistent with elevated neural ‘noise’ and altered timing and coordination of neural activity. This reduction was associated with lower verbal communication, suggesting the relevance of these neural measures for autistic youths’ communication abilities. It is important to note, however, that we did not find relationships with language skills measures with CELF.

Although we showed between-group differences in the neural measures, we identified similar age-related changes in the EEG metrics in both groups of participants. Periodic gamma power in response to speech stimuli increased with age, similar to previous findings (85). This age-related increase of gamma strength was attributed to developmental changes in E/I balance in local circuits, mostly by maturation of the inhibitory GABA system (85); this development starts early in neonate brain by switching from a depolarizing to hyperpolarizing action of GABA receptors, i.e., excitatory-to-inhibitory shift of GABA receptors (86). The decline in aperiodic exponent and offset observed in this study is also consistent with previous reports (45,74,87,88). These age-related declines may reflect the overall broadband change of background neural firing and may account for a normative redistribution of spectral power from lower to higher levels over development (45). However, it is still largely unexplored how age-related changes in these two types of neural activity – aperiodic power spectra and gamma-band oscillations – relate to each other. Future studies would benefit from combining these EEG measures with magnetic resonance spectroscopy to identify how concentrations of specific neurotransmitters (GABA vs. glutamate) and their age-related maturation are associated with functioning of EEG-based E/I balance metrics (87).

## Limitations

This work has some limitations that should be highlighted. First, although the study included a large sample size, most individuals with ASD were ‘high functioning’ with average or above average verbal skills. To generalize the findings and assess the validity of identified neural markers, it is necessary to include nonverbal or minimally verbal autistic individuals. Second, the study speculated on E/I balance and GABA/glutamate ratio based on non-invasive EEG measures. Future studies would benefit from combining these EEG measures with magnetic resonance spectroscopy to identify how concentration of specific neurotransmitters (GABA vs. glutamate) is related to EEG measures.

## Conclusion

The present study aimed to investigate a set of E/I balance measures derived from EEG speech perception task and their relation to clinical phenotype in a large group of youths with ASD. The main finding is that aperiodic component of spectral power was reduced in autistic youths compared to the TD group and this reduction was related to lower verbal communication skills in autistic youths. The findings suggested increased excitation and the reduction of broadband neural firing / spiking activity and, thus, higher ‘noisiness’ in the cortical systems responsible for speech perception (most likely, primary and secondary areas of the auditory cortex). This E/I imbalance pattern, therefore, may be relevant for monitoring language and communication skills in individuals with ASD across development. However, more studies addressing these metrics are needed to clarify their stability, validity, and generalizability.

## Declarations

### Ethics approval and consent to participate

The study was approved by the Yale Institutional Review Board, the UCLA Office of Human Research Protection Program, Boston Children’s Hospital Institutional Review Board, USC Office for the Protection of Research Subjects, and the University of Virginia Institutional Review Board for Health Sciences Research. All procedures performed were in accordance with the Declaration of Helsinki. All minor children provided verbal assent to participate in the study and were informed that they can withdraw from the study at any time during the experiment. A written consent form was obtained from a parent of each child participating in the study.

### Consent for publication

Not applicable

### Availability of data and materials

The behavioral and EEG data from the current study are available via the National Institute of Mental Health Data Archive Data Collection #2021.

## Acknowledgements

We wish to thank the families, parents, and children who participated in our study at our four data collection sites. The ACE GENDAAR Network additionally included contributions from: Katy Ankenman MSW, Elizabeth Aylward PhD, Veronica Kang PhD, Erin J. Libsack PhD, and Désirée Lussier-Lévesque PhD who were formerly at Seattle Children’s Research Institute; Sarah Corrigan MA and Waylon Howard PhD who are currently at Seattle Children’s Research Institute; Laura A. Edwards PhD and Jack Keller who were formerly at Boston Children’s Hospital; Rachael Tillman PhD who was formerly at Yale Child Study Center; Scott Huberty PhD who was formerly at UCLA; Zachary Jacokes who is currently at University of Virginia; Carinna Torgerson who is currently at USC; and Charles Nelson who is currently at Boston Children’s Hospital and Harvard Medical School. Additional important contribution was provided by Morgan Opdahl who is currently at UC Irvine.

## Authors’ contributions

VA, MS, EN, HB, RAB, SYB, MD, ARG, AJ, SJ, JCM, AN, JDVH, KAP, and SJW participated in project conceptualization and writing, including editing and final approval. MS, HB, SJ, AN, and SJW were involved in EEG data acquisition. VA, MS, HB, and SJW completed the analysis. VA and SJW drafted the manuscript.

## References

1. American Psychiatric Association. Diagnostic and statistical manual of mental disorders: DSM-5 (5th ed.). Washington, DC; London: American Psychiatric Publication, 2013.

2. Kjelgaard MM, Tager-Flusberg H. An investigation of language impairment in autism: Implications for genetic subgroups. Language and Cognitive Processes 2001; 16: 287–308.

3. Pickles A, Anderson DK, Lord C. Heterogeneity and plasticity in the development of language: a 17-year follow-up of children referred early for possible autism. The Journal of Child Psychology and Psychiatry 2014; 55: 1354–1362.

4. Tager-Flusberg H, Kasari K. Minimally Verbal School-Aged Children with Autism Spectrum Disorder: The Neglected End of the Spectrum. Autism Research 2013; 6: 468–478.

5. Tager-Flusberg H. Risk Factors Associated With Language in Autism Spectrum Disorder: Clues to Underlying Mechanisms. Journal of Speech, Language and Hearing Research 2016; 59: 143–154.

6. Crasto R, Folhas D, Pereira T, Lousada M. Clinical pathways from the perception of the first signs to the diagnosis of autism spectrum disorder in Portugal: A brief report of a survey of parents. Research in Autism Spectrum Disorders 2024; 114: 102382.

7. Miranda A, Berenguer C, Baixauli I, Rosello B. Childhood language skills as predictors of social, adaptive and behavior outcomes of adolescents with autism spectrum disorder. Research in Autism Spectrum Disorders 2023; 103: 102143.

8. Antoine MW, Langberg T, Schnepel P, Feldman DE. Increased Excitation-Inhibition Ratio Stabilizes Synapse and Circuit Excitability in Four Autism Mouse Models. Neuron 2019; 101(4): 648–661.

9. Gatto CL, Broadie K. Genetic controls balancing excitatory and inhibitory synaptogenesis in neurodevelopmental disorder models. Frontiers Synaptic Neuroscience 2010; 2: 4.

10. Levin R, Nelson CA. Inhibition-Based Biomarkers for Autism Spectrum Disorder. Neurotherapeutics 2015; 12: 546–552.

11. Rubenstein JLR, Merzenich MM. Model of autism: increased ratio of excitation / inhibition in key neural systems. Genes, Brain and Behavior 2003; 2: 255–267.

12. Sohal VS, Rubenstein JLR. Excitation-inhibition balance as a framework for investigating mechanisms of neuropsychiatric disorders. Molecular Psychiatry 2019; 24: 1248–1257.

13. Yizhar O, Fenno LE, Prigge M, Schneider F, Davidson TJ, O’Shea DJ, et al. Neocortical excitation / inhibition balance in information processing and social dysfunction. Nature 2011; 477: 171–178.

14. Contractor A, Ethell IM, Portera-Cailliau C. Cortical interneurons in autism. Nature Neuroscience 2021; 24: 1648–1659.

15. Tang X, Jaenisch R, Sur M. The role of GABAergic signalling in neurodevelopmental disorders. Nature Reviews Neuroscience 2021; 22: 290–307.

16. Chao HT, Chen H, Samaco RC, Xue M, Chahrour M, Yoo J, et al. Dysfunction in GABA signalling mediates autism-like stereotypies and Rett syndrome phenotypes. Nature 2010; 468(7321): 263–269.

17. Delattre V, La Mendola D, Meystre J, Markram H, Markram K. Nlgn4 knockout induces network hypo-excitability in juvenile mouse somatosensory cortex in vitro. Scientific Reports 2013; 3: 2897.

18. Dickinson A, Jones M, Milne E. Measuring neural excitation and inhibition in autism: Different approaches, different findings and different interpretations. Brain Research 2016; 1648(A): 277–289.

19. Arutiunian V, Arcara G, Buyanova I, Davydova E, Pereverzeva D, Sorokin A, et al. Neuromagnetic 40 Hz Auditory Steady-State Response in the left auditory cortex is related to language comprehension in children with Autism Spectrum Disorder. Progress in Neuropsychopharmacology and Biological Psychiatry 2023; 122: 110690.

20. Arutiunian V, Arcara G, Buyanova I, Fedorov M, Davydova E, Pereverzeva D, et al. Abnormalities in both stimulus-induced and baseline MEG alpha oscillations in the auditory cortex of children with Autism Spectrum Disorder. Brain Structure and Function 2024; 229:1225–1242.

21. Arutiunian V, Santhosh M, Neuhaus E, Borland H, Tompkins C, Bernier RA, et al. The relationship between gamma-band neural oscillations and language skills in youth with Autism Spectrum Disorder and their first-degree relatives. Molecular Autism 2024; 15: 19.

22. Bloy L, Shwayder K, Blaskey L, Roberts TPL, Embick D. A Spectrotemporal Correlate of Language Impairment in Autism Spectrum Disorder. Journal of Autism and Developmental Disorders 2019; 49: 3181–3190.

23. Plueckebaum H, Meyer L, Beck A-K, Menn KH. The developmental trajectory of functional excitation-inhibition balance relates to language abilities in autistic and allistic children. Autism Research 2023; 16(9): 1681–1692.

24. Wilkinson CL, Chung H, Dave A, Tager-Flusberg H, Nelson CA. Changes in Early Aperiodic EEG Activity Are Linked to Autism Diagnosis and Language Development in Infants With Family History of Autism. Autism Research 2025; 18: 1356–1368.

25. Wilkinson CL, Nelson CA. Increased aperiodic gamma power on young boys with Fragile X Syndrome is associated with better language ability. Molecular Autism 2021; 12: 17.

26. Agetsuma M, Hamm JP, Tao K, Fujisawa S, Yuste R. Parvalbumin-Positive Interneurons Regulate Neuronal Ensembles in Visual Cortex. Cerebral Cortex 2018; 28: 1831–1845.

27. Cardin JA, Carlén M, Meletis K, Knoblich U, Zhang F, Deisseroth K, Tsai L-H, Moore CI. Driving fast-spiking cells induces gamma rhythm and controls sensory responses. Nature 2009; 459: 663–668.

28. Carlén M, Meletis K, Siegle JH, Cardin JA, Futai K, Vierling-Claassen D, et al. A critical role for the NMDA receptors in parvalbumin interneurons for gamma rhythm induction and behavior. Molecular Psychiatry 2012; 17: 537–548.

29. Espinoza C, Guzman SJ, Zhang X, Jonas P. Parvalbumin+ interneurons obey unique connectivity rules and establish a powerful lateral-inhibition microcircuit in dentate gyrus. Nature Communication 2018; 9: 4605.

30. Ferguson BR, Gao W-J. PV Interneurons: Critical Regulators of E/I Balance for Prefrontal Cortex-Dependent Behavior and Psychiatric Disorders. Frontiers in Neural Circuits 2018; 12: 37.

31. Magueresse CL, Monyer H. GABAergic Interneurons Shape the Functional Maturation of the Cortex. Neuron 2013; 77: 388–405.

32. Gandal MJ, Edgar JC, Ehrlichman RS, Mehta M, Roberts TPL, Siegel SJ. Validating γ oscillations and delayed auditory responses as translational biomarkers of autism. Biological Psychiatry 2010; 68: 1100–1106.

33. Braeutigam S, Swithenby SJ, Bailey AJ. Contextual integration the unusual way: a magnetoencephalographic study of responses to semantic violation in individuals with autism spectrum disorders. European Journal of Neuroscience 2008; 27: 1026–1036.

34. McFadden KL, Hepburn S, Winterrowd E, Schmidt G, Rojas DC. Abnormalities in gamma-band responses to language stimuli in first-degree relatives of children with autism spectrum disorder: an MEG study. BMC Psychiatry 2012; 12: 213.

35. Ortiz-Mantilla S, Cantiani C, Shafer VL, Benasich AA. Minimally-verbal children with autism show deficits in theta and gamma oscillations during processing of semantically-related visual information. Scientific Reports 2019; 9: 5072.

36. Roberts TPL, Bloy L, Liu S, Ku M, Blaskey L, Jackel C. Magnetoencephalography Studies of the Envelope Following Response During Amplitude-Modulated Sweeps: Diminished Phase Synchrony in Autism Spectrum Disorder. Frontiers in Human Neuroscience 2021; 15: 787229.

37. Rojas DC, Maharajh K, Teale P, Rogers SJ. Reduced neural synchronization of gamma-band MEG oscillations in first-degree relatives of children with autism. BMC Psychiatry 2008; 8: 66.

38. Rojas DC, Teale PD, Maharajh K, Kronberg E, Youngpeter K, Wilson LB, et al. Transient and steady-state auditory gamma-band responses in first-degree relatives of people with autism spectrum disorder. Molecular Autism 2011; 2: 11.

39. Wang X, Delgado J, Marchesotti S, Kojovic N, Sperdin HF, Rihs TA, et al. Speech reception in young children with autism is selectively indexed by a neural oscillation coupling anomaly. Journal of Neuroscience 2023; 43: 6779–6795.

40. Wilson TW, Rojas DC, Reite ML, Teale PD, Rogers SJ. Children and Adolescents with Autism Exhibit Reduced MEG Steady-State Gamma Responses. Biological Psychiatry 2007; 62: 192–197.

41. Romeo RR, Choi B, Gabard-Durnam LJ, Wilkinson CL, Levin AR, Rowe ML, et al. Parental Language Input Predicts Neuroscillatory Patterns Associated with Language Development in Toddlers at Risk of Autism. Journal of Autism and Developmental Disorders, 2022; 52: 2717–2731.

42. Wilkinson CL, Levin AR, Gabard-Durnam LJ, Tager-Flusberg H, Nelson CA. Reduced frontal gamma power at 24 months is associated with better expressive language in toddlers at risk for autism. Autism Research 2019; 12: 1211–1224.

43. Donoghue T, Haller M, Peterson EJ, Varma P, Sebastian P, Gao R, et al. Parameterizing neural power spectra into periodic and aperiodic components. Nature Neuroscience 2020; 23: 1655–1665.

44. Levin AR, Naples AJ, Scheffler AW, Webb SJ, Shic F, Sugar CA, et al. Day-to-Day Test-Retest Reliability of EEG Profiles in Children With Autism Spectrum Disorder and Typical Development. Frontiers in Integrative Neuroscience 2020; 14: 21.

45. Ostlund B, Donoghue T, Anaya B, Gunther KE, Karalunas SL, Voytek B, Pérez-Edgar KE. Spectral parameterization for studying neurodevelopment: How and why. Developmental Cognitive Neuroscience 2022; 54: 101073.

46. Gao R, Peterson EJ, Voytek B. Inferring synaptic excitation/inhibition balance from field potentials. NeuroImage 2017; 158: 70–78.

47. Manning JR, Jacobs J, Fried I, Kahana MJ. Broadband shifts in local field potential power spectra are correlated with single-neuron spiking in humans. Journal of Neuroscience 2009; 29(43): 13613–13620.

48. Miller KJ. Broadband Spectral Change: Evidence for a Macroscale Correlate of Population Firing Rate? Journal of Neuroscience 2010; 30(19): 6477–6479.

49. American Psychiatric Association. Diagnostic and statistical manual of mental disorders: DSM-IV-TR. Washington, DC: Author, 2000.

50. Lord C, Rutter M, DiLavore PC, Risi S, Gotham K, Bishop S. Autism Diagnostic Observation Schedule, second edition (ADOS-2) manual (part I): modules 1–4. Torrance, CA: Western Psychological Services, 2012.

51. Lord C, Rutter M, Le Couteur A. Autism Diagnostic Interview—Revised: A revised version of a diagnostic interview for caregivers of individuals with possible pervasive developmental disorders. Journal of Autism and Developmental Disorders 1994; 24(5): 659–685.

52. Sparrow S, Cicchetti D, Balla D. Vineland Adaptive Behavior Scales, second edition (Vineland-II). Circle Pines, MN: American Guidance Service, 2005.

53. Elliott CD. Differential Ability Scales, second edition. San Antonio, TX: The Psychological Corporation, 2007.

54. Semel E, Wiig EH, Secord WA. Clinical evaluation of language fundamentals, fourth edition (CELF-4). Toronto: The Psychological Corporation / A Harcourt Assessment Company, 2003.

55. Constantino JN. Social Responsiveness Scale, second edition (SRS-2). Torrance, CA: Western Psychological Services, 2012.

56. Rutter ML, Bailey A, Lord C. Social Communication Questionnaire. Torrance, CA. Western Psychological Services, 2003.

57. Arnett AB, Hudac CM, DesChamps TD, Cairney BM, Gerdts J, Wallace AS, et al. Auditory perception is associated with implicit language learning and receptive language ability in autism spectrum disorder. Brain and Language 2018; 187: 1–8.

58. Liu J, Tsang T, Ponting C, Jackson L, Jeste SS, Bookheimer SY, Dapretto M. Lack of neural evidence for implicit language learning in 9-month-old infants at high risk for developing autism. Developmental Science 2021; 24: e13078.

59. McNealy K, Mazziotta JC, Dapretto M. Age and experience shape developmental changes in the neural basis of language-related learning. Developmental Science 2011; 14: 1261–1282.

60. McNealy K, Mazziotta JC, Dapretto M. The Neural Basis of Speech Parsing in Children and Adults. Developmental Science 2010; 13: 385–406.

61. Saffran JR, Aslin RN, Newport EL. Statistical learning by 8-months-old infants. Science 1996; 274: 1926–1928.

62. Scott-Van Zeeland AA, McNealy K, Wang AT, Sigman M, Bookheimer SY, Dapretto M. No neural evidence of statistical learning during exposure to artificial languages in children with autism spectrum disorders. Biological Psychiatry 2010; 68: 345–351.

63. Levin AR, Méndez Leal AS, Gabard-Durnam LJ, O’Leary HM. BEAPP: The Batch Electroencephalography Automated Processing Platform. Frontiers in Neuroscience 2018; 12: 513.

64. Gabard-Durnam LJ, Méndez Leal AS, Wilkinson CL, Levin AR. The Harvard Automated Processing Pipeline for Electroencephalography (HAPPE): Standardized Processing Software for Developmental and High-Artifact Data. Frontiers in Neuroscience 2018; 12: 97.

65. R Core Team. R: A Language and Environment for Statistical Computing. R Foundation for Statistical Computing. Vienna, 2019. URL. https://www.R-project.org/

66. Bates D, Mächler M, Bolker BM, Walker SC. Fitting linear mixed-effects models using *lme4*. Journal of Statistical Software 2015; 67: 1–48.

67. Lüdecke D. sjPlot: Data Visualization for Statistics in Social Science. R package version 2.8.4, 2020. URL: https://CRAN.R-project.org/package=sjPlot

68. Wickham H. ggplot 2: Elegant Graphics for Data Analysis. New York: Springer-Verlag, 2016.

69. Hunter JD. Matplotlib: a 2D graphics environment. Computing in Science and Engineering 2007; 9: 90–95.

70. Freeman WJ, Zhai J. Simulated power spectral density (PSD) of background electrocorticogram (ECoG). Cognitive Neurodynamics 2009; 3: 97–103.

71. Hancock R, Pugh KR, Hoeft F. Neural Noise Hypothesis of Developmental Dyslexia. Trends in Cognitive Sciences 2017; 21(6): 434–448.

72. Pi Y, Pscherer C, Muckschel M, Colzato L, Hommel B, Beste C. Metacontrol-related aperiodic neural activity decreases but strategic adjustment thereof increases from childhood to adulthood. Scientific Reports 2025; 15: 18349.

73. Voytek B, Knight RT. Dynamic Network Communication as a Unifying Neural Basis for Cognition, Development, Aging, and Disease. Biological Psychiatry 2015; 77(12): 1089–1097.

74. Voytek B, Kramer MA, Case J, Lepage KQ, Tempesta ZR, Knight RT, Gazzaley A. Age-Related Changes in 1/f Neural Electrophysiological Noise. The Journal of Neuroscience 2015; 35: 13257–13265.

75. Parks LK, Hill DE, Thoma RJ, Euler MJ, Lewine JD, Yeo RA. Neural correlates of communication skill and symptom severity in autism: A voxel-based morphometry study. Research in Autism Spectrum Disorders 2009; 3: 444–454.

76. Wataru S, Shota U. The atypical social brain network in autism: advances in structural and functional MRI studies. Current Opinion in Neurology 2019; 32: 617–621.

77. Giraud A-L, Kleinschmidt A, Poeppel D, Lund TE, Frackwiak RSJ, Laufs H. Endogenous Cortical Rhythms Determine Cerebral Specialization for Speech Perception and Production. Neuron 2007; 56: 1127–1134.

78. Giraud A-L, Poeppel D. Cortical oscillations and speech processing: emerging computational principles and operations. Nature Neuroscience 2012; 15: 511–517.

79. Hämäläinen JA, Rupp A, Soltész F, Szücs D, Goswami U. Reduced phase locking to slow amplitude modulation in adults with dyslexia: An MEG study. NeuroImage 2012; 59: 2952–2961.

80. Moerel M, De Martino F, Formisano E. Processing of Natural Sounds in Human Auditory Cortex: Tonotopy, Spectral Tuning, and Relation to Voice Sensitivity. The Journal of Neuroscience 2012; 32(41): 14205–14216.

81. Moon J, Orlandi S, Chau T. A comparison and classification of oscillatory characteristics in speech perception and covert speech. Brain Research 2022; 1781: 147778.

82. Poeppel D, Assaneo MF. Speech rhythms and their neural foundations. Nature Reviews Neuroscience 2020; 21: 322–334.

83. Poeppel D. The analysis of speech in different temporal integration windows: cerebral lateralization as ‘asymmetric sampling in time’. Speech Communication 2003; 41: 245–255.

84. Zhang M, Zhu J, Jiang Y, Chang T, Fu Y, Wu K, et al. Excitation-inhibition imbalance in the auditory cortex causes sound recognition impairment in noisy environment in hidden hearing loss mice. Brain Research Bulletin 2025; 226: 111371.

85. Cho RY, Walker CP, Polizzotto NR, Wozny TA, Fissell C, Chen C-MA, Lewis DA. Development of Sensory Gamma Oscillations and Cross-Frequency Coupling from Childhood to Early Adulthood. Cerebral Cortex 2015; 25: 1509–1518.

86. Ben-Ari I. The GABA excitatory/inhibitory developmental sequence: A personal journey. Neuroscience 2014; 279: 187–219.

87. Hill AT, Clark GM, Bigelow FJ, Lum JAG, Enticott PG. Periodic and aperiodic neural activity displays age-dependent changes across early-to-middle childhood. Developmental Cognitive Neuroscience 2022; 54, 101076.

88. Merkin A, Sghirripa S, Graetz L, Smith AE, Hordacre B, Harris R, et al. Do age-related differences in aperiodic neural activity explain differences in resting EEG alpha? Neurobiology of Aging 2023; 121, 78–87.

